# GOLF: A Generative AI Framework for Pathogenicity Prediction of Myocilin OLF Variants

**DOI:** 10.1101/2025.06.17.660210

**Authors:** Thomas Walton, Darin Tsui, Lauren Fogel, Dustin J. E. Huard, Rafael Siqueira Chagas, Raquel L. Lieberman, Amirali Aghazadeh

## Abstract

Missense mutations in the *MYOC* gene, particularly those affecting the olfactomedin (OLF) domain of the myocilin protein, can be causal for open-angle glaucoma—a leading cause of irreversible blindness. However, predicting the pathogenicity of these mutations remains challenging due to the complex effects of toxic gain-of-function variants and the scarcity of labeled clinical data. Herein, we present GOLF, a generative AI framework for assessing and explaining the pathogenicity of OLF domain variants. GOLF collects and curates a comprehensive dataset of OLF homologs and trains generative models that predict the effect of monoallelic missense mutations. While these models exhibit diverse predictive behaviors, they collectively achieve accurate classification of known pathogenic and benign variants. To interpret their decision mechanisms, GOLF uses a sparse autoencoder (SAE) that reveals the underlying biochemical features exploited by the generative models to predict variant effects. GOLF enables accurate evaluation of disease-causing mutations, supporting early genetic risk stratification for glaucoma and facilitating interpretable investigations into the molecular basis of pathogenic variants.

## 1 Introduction

Understanding how genetic variation gives rise to disease phenotypes is essential for both advancing our knowledge of human genomics and enabling personalized clinical care. However, predicting the consequences of coding mutations, especially missense mutations, remains a major challenge. While loss-of-function mutations are often more straightforward to predict, missense mutations that result in toxic gain-of-function effects are considerably more difficult to characterize due to their subtle, context-dependent impact on protein behavior and cellular function^1^. In this work, we focus on a particularly important case: monoallelic missense mutations in the *MYOC* gene, which encodes the myocilin protein. Mutations in *MYOC*, especially those affecting its olfactomedin (OLF) domain, can lead to myocilin-associated open-angle glaucoma (OAG), a prevalent and irreversible form of blindness ^2–4^. OAG affects over 70 million people worldwide, with *MYOC* -linked cases estimated to account for approximately 3 million of these ^4–7^. The disease typically progresses without overt symptoms until permanent vision loss occurs, underscoring the critical need for early genetic risk stratification ^8^.

Disease-causing mutations in *MYOC* are concentrated within its olfactomedin (OLF) domain, a highly conserved region critical to myocilin’s structure and function ^2^. As a result, accurately assessing the pathogenicity of OLF domain variants is essential for interpreting the clinical significance of population-level *MYOC* mutations in glaucoma. However, doing so remains a challenge: experimental assays are time-consuming, and molecular simulations demand substantial computational resources. Current clinical evaluation often relies on detailed phenotypic data and multigenerational inheritance patterns ^9^, which are rarely available at the population scale.

With the increasing adoption of large-scale genotyping, a growing number of *MYOC* missense variants have been cataloged in public databases such as gnomAD^10^. Yet, most of these variants lack corresponding clinical annotations or familial segregation data, leaving their pathogenicity uncertain. This gap highlights the urgent need for computational approaches that can robustly assess the functional impact of OLF domain mutations in the absence of extensive clinical context. Generative models trained on evolutionary sequence data offer a principled approach for uncovering the sequence-level dependencies that determine whether a mutation is likely to be deleterious or tolerated. By leveraging natural variation observed across homologous protein sequences, typically via multiple sequence alignments (MSAs), these models learn statistical patterns that enable quantitative scoring of mutations. Likelihood-based predictions from such models have shown strong agreement with deep mutational scanning data and remain effective even when labeled clinical data are sparse or experimental throughput is limited ^11–14^. Critically, generative models can identify mutations that are functionally disruptive, even when the precise molecular function of the protein remains unknown. This is particularly advantageous for myocilin, whose biological role has not yet been fully elucidated ^7^. Moreover, recent advances in model interpretability allow extraction of biochemically meaningful features from generative models, shedding light on residue-level interactions and structural motifs that influence variant effects ^15–19^.

Herein, we hypothesize that generative models can learn the distribution of evolutionary OLF sequences and thereby predict the pathogenicity of novel missense mutations. To test this hypothesis, we introduce GOLF, a generative AI framework for variant effect prediction in the OLF domain of *MYOC* (Figure 1). GOLF collects a dataset of OLF homologs from UniRef100^20^, curates the sequences to construct diverse MSA data spanning multiple taxa, and trains a set of generative models to learn the distribution of OLF sequence variation. These models are then used to score all possible monoallelic missense mutations in the OLF domain. Finally, GOLF extracts and analyzes key features used in model predictions, enabling protein-wide interpretation of variant effects and supporting comprehensive analysis of pathogenicity. To summarize the contributions:

**Figure 1:**
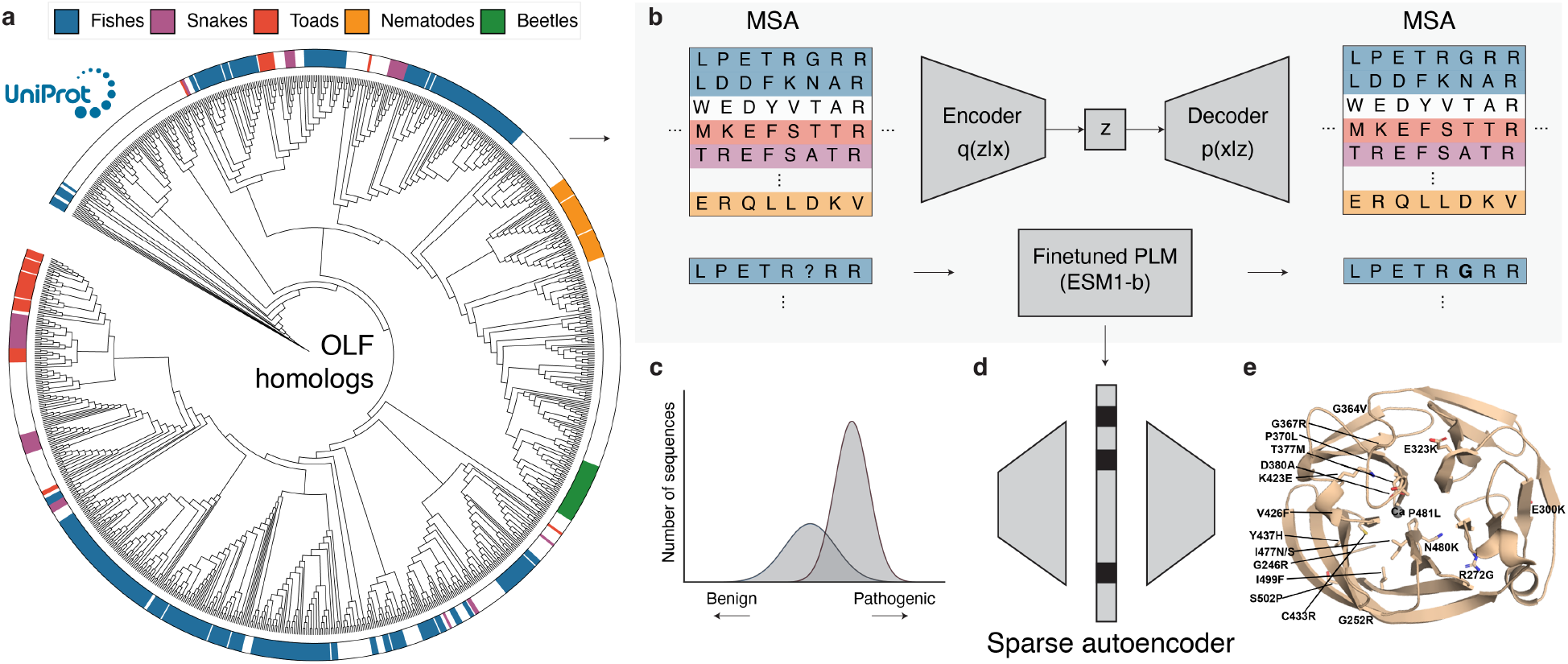
Schematic of GOLF, a generative AI framework for modeling and explaining missense mutations in the OLF domain of the myocilin protein. **a**, Phylogenetic tree of OLF homologs, collected from UniRef100. OLF homologs originate from a diverse set of 73 unique taxonomic groups. **b**, Multiple sequence alignments derived from a phylogenetic analysis encode evolutionary constraints of OLF, which can be used to train generative models to predict variant effect. **c**, Gaussian mixture models trained on the score from the generative models categorize mutations as pathogenic or benign. **d**, A sparse autoencoder (SAE) identifies key factors used by generative models to predict variant effects. **e**, 3D structure of the OLF domain, annotated with a subset of known pathogenic mutations. Pathogenic missense variants are distributed throughout the domain, highlighting the need for evolutionary context to accurately characterize their effects.

- We develop GOLF (<u>G</u>enerative <u>OLF</u>), a generative AI framework for predicting and explaining variant effects of missense mutations in the OLF domain of the myocilin protein.^1^
- We construct a curated dataset of OLF homologs spanning 73 taxonomic groups and perform a phylogenetic analysis to uncover the evolutionary origin and diversity of these sequences.
- We train two generative models, a variational autoencoder based on the EVE model ^11^ and a fine-tuned ESM-1b transfomer^21^, to learn the distribution of evolutionary OLF sequences.
- We collect a set of OLF missense variants with known pathogenicity labels from ClinVar and other published resources, and demonstrate that the variational autoencoder in GOLF classifies pathogenic and benign variants with over 96% accuracy.
- We train a sparse autoencoder (SAE) to extract learned features from generative models, finding that they use biochemically relevant rules, such as hydrophobicity and polarity, to infer pathogenicity.

## 2 The GOLF Framework

In this section, we describe the GOLF framework, which comprises three main components: (1) data collection and visualization, (2) model training and evaluation, and (3) feature interpretation.

### 2.1 Data Collection and Visualization

#### Gathering OLF homologs

The first step in GOLF is to gather homologous sequences of OLF proteins, which encapsulate the evolutionary context necessary for downstream analyses (Figure 1a). Following the EVCouplings framework ^22^, GOLF finds the MSA of OLF by performing five iterations of jackhmmer^23^ against the UniRef100 database ^20^. This initial search returned 40,026 sequences, which are further filtered to 11,867 by removing duplicates and dropping sequences shorter than 200 residues. To ensure phylogenetic diversity, we cluster the filtered set at 95% sequence identity and 80% coverage using MMSeqs2^24^, yielding 4,119 clusters. Finally, we sample one representative from each cluster to assemble a training dataset of OLF homologs.

#### Phylogenetic analysis

To better understand the evolutionary diversity of OLF, we visualize the phylogenetic relationships over a subset of homologs. We cluster our filtered dataset of 11,867 sequences to 80% sequence identity and sample one representative from each cluster, yielding 957 sequences. We select this lower identity threshold (as opposed to 95%) to ensure computational feasibility and visual clarity, as a lower sequence identity results in fewer representative sequences. We then compute the phylogenetic tree of these sequences using IQ-TREE^25,26^ and visualize them using the Interactive Tree Of Life (iTOL) web server^27^, resulting in 73 taxonomic groups.

### 2.2 Models, Training, and Evaluation

GOLF trains a set of generative models to learn the evolutionary constraints encoded in OLF sequence variation (Figure 1b,c). These models are designed to assign plausibility scores to candidate missense mutations, enabling both pathogenicity prediction and biochemical interpretation. Below, we describe the architecture, training, and evaluation for each model used within GOLF.

#### EVE

The Evolutionary model of Variant Effect (EVE) ^11^ is a Bayesian variational autoencoder (VAE) ^28^ that models the distribution of homologous protein sequences in MSA. This model enables unsupervised prediction of variant effects by learning the distribution of protein sequences that capture evolutionary constraints. EVE consists of an encoder and a decoder network, wherein the encoder maps the input sequence *x* to a lower-dimensional latent variable vector *z*, governed by a variational posterior *q*(*z* | *x*), while the decoder parameterizes the likelihood *p*(*x* | *z*), reconstructing sequences from the latent variables. To train the VAE, the evidence lower bound (ELBO) is minimized, balancing a reconstruction likelihood term and a Kullback–Leibler (KL) divergence regularizer, which encourages the model to learn a compact latent representation that reflects evolutionary structure. After training, the likelihoods of wild-type and mutant sequences will be computed from the decoder. EVE defines an evolutionary index for a mutation as the log-likelihood ratio between the mutant and the wild-type sequence: 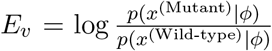, where *ϕ* denotes the learned parameters of the VAE^11^. *E*_*v*_ quantifies the degree to which a mutation deviates from the learned evolutionary distribution. These evolutionary indices are then clustered using a two-component Gaussian mixture model (GMM) to classify variants as benign or pathogenic. EVE also outputs an uncertainty score derived from its Bayesian decoder, which can be used to flag ambiguous predictions. GOLF reports predictions using the 100% retained class label (i.e., no variants withheld) to ensure complete coverage across the variants.

GOLF trains EVE on the final dataset of 4,119 OLF homologs. We evaluated a range of latent dimensionalities and found that a latent dimension of 40 provided the best performance. Larger latent spaces tended to bias the model toward predicting pathogenicity, while smaller ones favored benign classifications. The training was performed for 200k training steps, after which we observed signs of overfitting. To assess the impact of hyperparameter choices, we held out 20% of the sequences as a test set and monitored their ELBO after training. Larger latent dimensions more effectively minimized KL divergence but at the cost of increased reconstruction error. Conversely, smaller latent dimensions achieved better reconstruction but resulted in less regularized latent spaces. Across all settings, ELBO values were comparable, reflecting a tradeoff between latent space regularization and sequence reconstruction fidelity.

#### ESM-1b

ESM-1b^21^ is a large transformer-based protein language model (pLM) pretrained on 250 million protein sequences spanning 19,000 distinct protein families. To adapt ESM-1b to the OLF domain for GOLF, we fine-tuned the model on our curated dataset of 4,119 OLF homologs. Specifically, we fine-tuned the last three layers and froze the remaining parameters to prevent overfitting and ensure training stability, following standard fine-tuning practices for domain-specific datasets ^29,30^. Fine-tuning was performed for 30 epochs using the masked language modeling (MLM) objective, with a learning rate of 10^−5^ and a cosine decay schedule. Hyperparameters were selected using a 10% validation split. After fine-tuning, we computed the evolutionary indices for each mutation, following the same procedure as in EVE. We then applied a two-component GMM to these scores to classify mutations as pathogenic or benign.

#### Benchmarking with AlphaMissense

AlphaMissense ^31^ is a large-scale, structure-informed model for predicting the pathogenicity of human missense variants. It builds on AlphaFold2^32^ by incorporating both evolutionary information and structural context derived from AlphaFold-predicted protein models. During training, common missense variants observed in human and primate populations are treated as benign, while rare or unobserved variants are labeled as pathogenic. Rather than generating new structures for each mutation, AlphaMissense uses precomputed AlphaFold2 predictions of the wild-type structure and evaluates how substitution is likely to affect local structural confidence and residue environment. The model outputs a pathogenicity score between 0 and 1. To ensure a fair comparison with our methods, we thresholded AlphaMissense scores at 0.5: variants scoring above this threshold were classified as pathogenic, and those below as benign. Ambiguous predictions were treated the same way to maintain consistency across evaluations.

#### Interpreting features

The final component of GOLF leverages a sparse autoencoder (SAE) to extract interpretable features associated with pathogenicity (Figure 1d). Recent studies have shown that generative models trained on evolutionary sequences implicitly capture structural and biochemical constraints from sequence variations alone ^16,33,34^. However, such models largely remain inaccessible for human interpretation. To address this, SAEs learn compressed and interpretable latent representations of model embeddings by promoting sparsity in the latent space, where individual latent variables activate in response to specific input features ^35,36^. These activations often correspond to biologically meaningful patterns, such as structural motifs or biochemical properties ^18,19^. By analyzing these latent variables, we gain insight into the features generative models rely on when predicting variant effects.

We employ an SAE^19^ trained on frozen embeddings from ESM2^37^. To identify latent variables associated with pathogenicity, we use linear probes to predict EVE scores for single missense mutations. Mean-pooled embeddings from ESM2 are passed through the pretrained SAE to produce sparse latent representations, which are then used as inputs to a ridge regression model. Regularization strength is selected via grid search on a validation set. We feed the wild-type OLF domain sequence into the SAE to obtain latent activations and focus on visualizing the latent variables associated with benign and pathogenic predictions. We project the active residues for each latent onto the OLF structure (PDB ID 4WXQ^38^, residues 244–504) to demonstrate where these features are concentrated and infer potential biochemical relevance. Full details are provided in Appendix A, and the results are presented in the next section.

## 3 Results

### Phylogenetic analysis reveals taxonomic diversity

Figure 1a shows the phylogenetic tree constructed from a representative subset of OLF homologs. Notably, we observed that a majority of sequences were not derived from humans, with a substantial proportion of approximately 40% of sequences derived from fishes. Surprisingly, 5% of sequences originate from nematodes, a phylum of simple invertebrates that lack visual systems entirely. Despite their anatomical divergence from humans, nematode OLF domains remain highly conserved at the sequence level. These findings highlight the evolutionary conservation of the OLF domain and its widespread presence across diverse taxa. This diversity provides a rich foundation for learning generalizable evolutionary constraints, which is critical for accurately predicting the pathogenicity of novel missense mutations (see Appendix B for the full list of taxonomic groups).

### EVE accurately classifies OLF variants

Table 1 details the predictive performance of EVE compared to fine-tuned ESM-1b and AlphaMissense on a set of 32 OLF mutants with generally accepted pathogenicity labels (see Appendix C and Figure 2a). EVE achieved the highest accuracy among all models, correctly identifying every pathogenic variant and misclassifying only one benign mutation. In contrast, both ESM-1b and AlphaMissense incorrectly labeled 10 pathogenic variants as benign. Notably, both ESM-1b and AlphaMissense demonstrated reduced performance for mutations occurring between residues 246–300, but performed better on pathogenic variants located between residues 380–480. The reduced performance of ESM-1b and AlphaMissense for mutations between residues 246-300 is likely due to limited clinical and experimental data for this region, restricting the models’ ability to make accurate predictions ^9^. Additionally, this segment lies near the interface between blades E and A of the *β*-propeller, a region known for structural irregularities, which may further contribute to the difficulty in categorizing variants ^39^. In contrast, the improved performance observed for mutations between residues 380-480 may reflect the proximity of this region to functionally important features such as the metal-binding center and the disulfide clasp. These elements are critical for proper folding and stability ^38^, making mutations in this region more disruptive and, consequently, better characterized in available datasets.

**Table 1:** Performance of variant effect prediction methods on a held-out set of 32 variants (7 benign, 25 pathogenic). A full set of variant predictions is presented in Figure 2.

| Metric | ESM-1b | AlphaMissense | EVE |
| --- | --- | --- | --- |
| Accuracy (overall) | 68.8% | 68.8% | 96.9% |
| Accuracy (benign) | 100.0% (7/7) | 100.0% (7/7) | 85.7% (6/7) |
| Accuracy (pathogenic) | 60.0% (15/25) | 60.0% (15/25) | 100.0% (25/25) |

**Figure 2:**
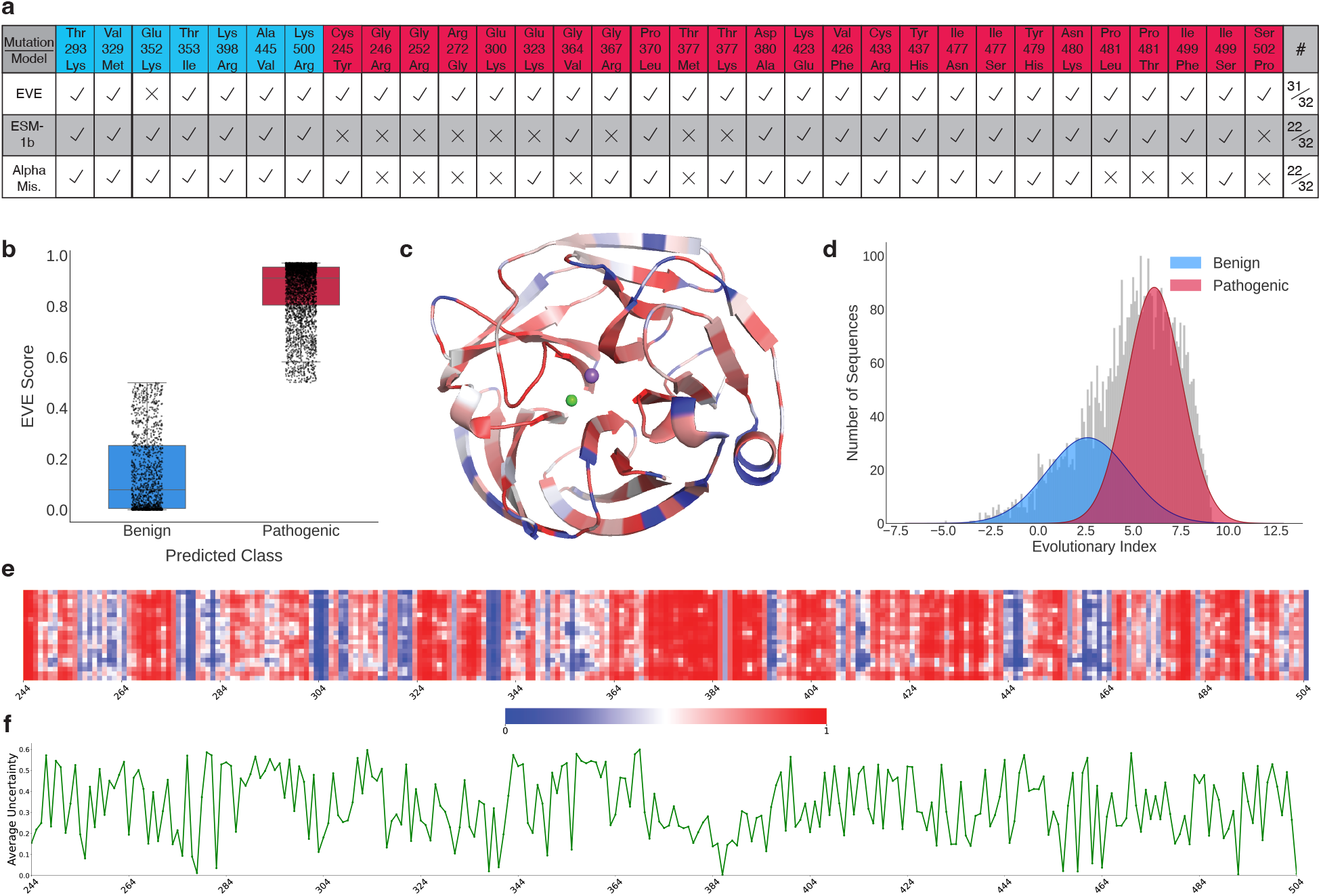
Overview of variant effect predictions on the OLF domain. **a**, Comparison of predictions across 32 single-residue OLF mutations with known labels. Blue labels indicate a benign mutation, whereas red indicates pathogenic. **b**, Distribution of EVE scores. Scores closer to 0 indicate a benign prediction, and scores closer to 1 indicate a pathogenic prediction. **c**, Structure of the OLF domain colored according to the average predicted variant effect at each position. Regions shaded in red are predicted by EVE to be associated with pathogenicity upon mutation, whereas blue regions indicate positions more likely to tolerate or benefit from variation. Lighter colors reflect positions near the boundary between predicted pathogenic and benign effects. **d**, Distribution of evolutionary indices predicted by EVE. The benign and pathogenic distributions are determined by a two-component Gaussian Mixture Model (GMM). The distribution of mutant OLF sequences is indicated in gray. **e**, Heatmap of mutational tolerance across all positions in the OLF domain (residues 244-504 of myocilin). Each column corresponds to a sequence position, and each row represents one of the 20 residues. For every position, the EVE score is computed for all 19 possible single-residue mutations and visualized as a matrix, where each cell reflects the predicted pathogenicity of a specific substitution. Lighter regions denote ambiguous predictions or the wild-type residue at that position. **f**, Average uncertainty per residue in EVE. Uncertainty is determined from the decoder of EVE, ranging on a scale of 0 (high confidence) to 1 (high uncertainty).

EVE’s evolutionary indices exhibited a bimodal distribution, with most variants clustering near 0 or 1, corresponding to predicted benign and pathogenic effects, respectively (Figure 2b). This separation indicates that EVE effectively learned between tolerated and deleterious mutations. Figure 2c illustrates the average mutational effect mapped to the structure of the OLF domain. Across the full mutational landscape of the OLF domain, EVE predicted 3,503 out of 4,959 possible missense variants as pathogenic, with the remainder classified as benign (Figure 2d). This skew toward pathogenic predictions reflects the model’s sensitivity to deviations from conserved sequence patterns. A summary of predicted pathogenicity of all possible missense mutations across each residue in the OLF domain is presented in Figure 2e. We identified several regions with heightened sensitivity to mutation, particularly residues 266–290, 324–334, and 363–394. An exception was residue 386, which remained tolerant to most substitutions despite being located in a generally sensitive region. EVE’s Bayesian decoder provides an uncertainty score for each prediction, enabling position-wise estimation of confidence. Figure 2f displays the average uncertainty across OLF. We observed low uncertainty in regions where scores were close to 0 or 1, with spikes in uncertainty occurring where EVE scores deviated from these extremes.

### Effect of ensembling

We observed that the predictive performance of EVE was sensitive to initialization. To mitigate this, we trained an ensemble of five independently initialized EVE models, each using the full set of OLF sequences and identical hyperparameter configurations. Following training, we computed the evolutionary indices from each model and averaged them. We clustered the resulting indices, resulting in pathogenic and benign labels for each single-residue mutation to OLF. We found that ensembling increased performance by 4% compared to the average performance of a single EVE model when predicting known mutation labels.

### Generative models implicitly learn biochemical rules underlying the variant effects

Our analysis of the top SAE latent variables reveals a clear biochemical distinction: SAE features most strongly associated with benign predictions exhibited high activations for aliphatic hydrophobic residues, such as valine (V) and isoleucine (I) (Figure 3a). In contrast, features most predictive of pathogenicity exhibited high activations for polar or aromatic residues, such as threonine (T) and phenylalanine (F) (Figure 3b). This mirrors our existing knowledge of the OLF domain, where hydrophobic residues are known to stabilize the *β*-propeller through tight packing, whereas polar residues are more likely to disrupt structural integrity and promote misfolding ^38^. Strikingly, these patterns emerge without explicit supervision, indicating that generative models capture biochemical properties from sequence alone.

**Figure 3:**
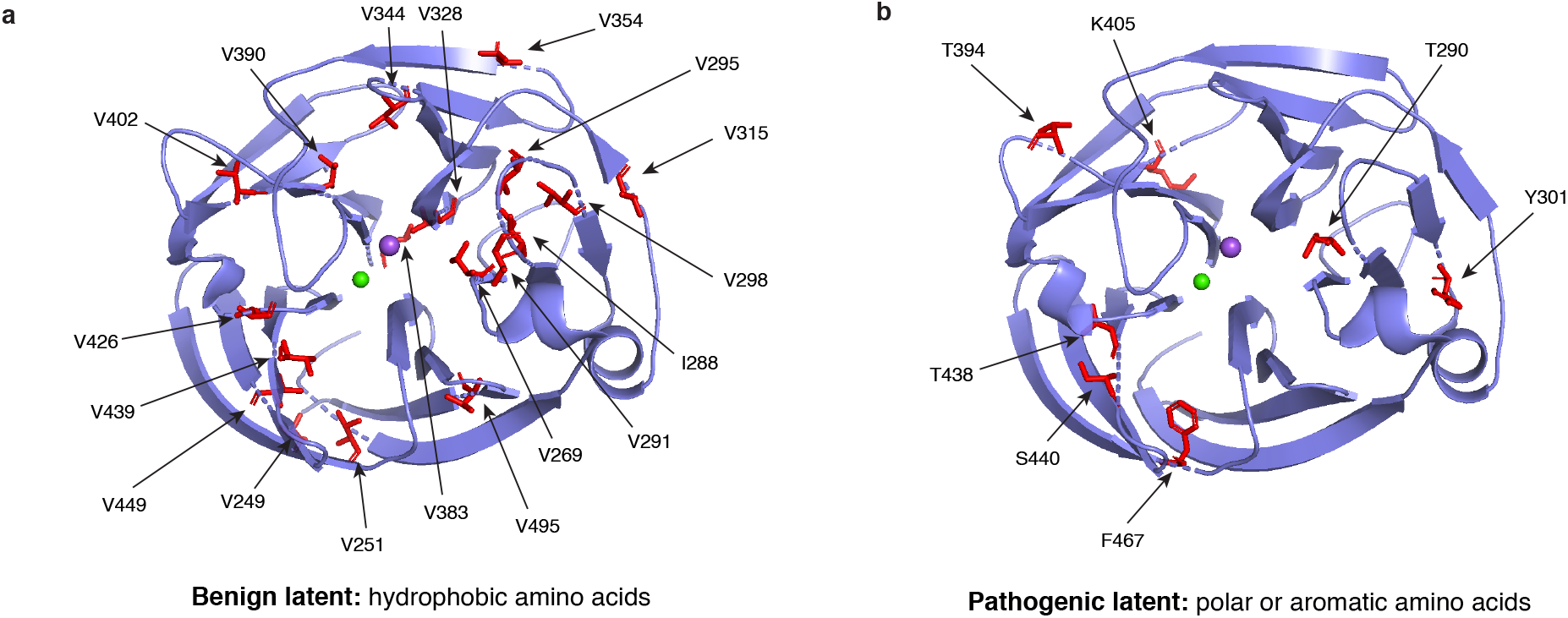
Mechanistic interpretability of the disease prediction models. Visualization of the latent variables associated with **a**, benign and **b**, pathogenic predictions. Benign latent variables favored hydrophobic residues, while pathogenic latent variables favored polar or aromatic residues.

## 4 Conclusion and Discussion

In this study, we introduced GOLF, a generative modeling framework developed to predict the distribution of pathogenic and benign variants across the OLF domain of myocilin. GOLF achieved high predictive accuracy, correctly classifying all but one of the labeled variants. By accurately predicting the pathogenicity of OLF missense variants, our generative modeling approach addresses the critical need for rapid screening of *MYOC* mutations, facilitating early risk stratification in glaucoma screening.

Beyond predictive performance, GOLF provides interpretable insights into the biochemical determinants of variant effects. Using a sparse autoencoder, we found that latent features align with established biochemical properties, such as hydrophobicity and polarity, that influence mutational tolerance. These results suggest that generative models can implicitly learn fundamental principles of protein biochemistry, capturing residue-level interactions and structural constraints without explicit supervision. We anticipate that GOLF will serve as a foundation for future studies aiming to uncover the mechanistic basis of disease and extend interpretable variant effect prediction to other clinically relevant protein domains.

### Limitations and future work

While GOLF demonstrates strong performance in predicting the pathogenicity of OLF variants, its validation is limited by the small number of labeled mutations currently available. As more clinically annotated variants become accessible, we anticipate that GOLF’s predictive power and calibration will continue to improve. A current limitation of generative models, including GOLF, is their inability to distinguish between distinct functional outcomes, specifically, gain-of-function versus loss-of-function mutations. For instance, the Glu352Lys variant, which is more prevalent in African American populations who are also at a generally elevated glaucoma risk ^40^, may contribute to disease through a subtle gain-of-function mechanism or a loss of function. Although this variant was classified as benign *in vitro* ^4,9^, and similarly predicted as benign by AlphaMissense and ESM-1b, it was predicted as pathogenic by EVE. This discrepancy suggests that EVE may capture aspects of pathogenicity that are overlooked by other models. Our sparse autoencoder analysis provides interpretable insights by linking latent dimensions to biochemical properties such as hydrophobicity and polarity. While these patterns offer promising hypotheses about the biochemical basis of pathogenicity, experimental validation is essential to confirm whether the implicated residues and features reflect causal mechanisms. Designing targeted experiments guided by latent space features represents an exciting future direction that could bridge the gap between model interpretability and biological discovery.

## Acknowledgments

This work was supported by the Parker H. Petit Institute for Bioengineering and Biosciences (IBB) interdisciplinary seed grant, the National Science Foundation (NSF) Graduate Research Fellowship Program (GRFP), the Georgia Institute of Technology start-up funds (A.A.), and NIH R01EY021205 (R.L.L.).

## Appendix

### A Sparse Autoencoder Details

#### TopK SAEs

We use sparse autoencoders (SAEs) with TopK activation ^35^ to enforce sparsity in the latent space. The TopK function retains only the *k* largest activations and sets all others to zero. Given an input vector *x* ∈ ℝ^*n*^, the encoder maps *x* to a latent representation *z* ∈ ℝ^*d*^ via:

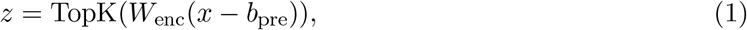

where *W*_enc_ ∈ ℝ^*d×n*^ is a learnable weight matrix and *b*_pre_ ∈ ℝ^*n*^ is a bias term. The decoder reconstructs the input as:

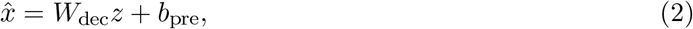

with *W*_dec_ ∈ ℝ^*n×d*^. The SAE is trained to minimize the mean squared error between *x* and its reconstruction 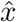:

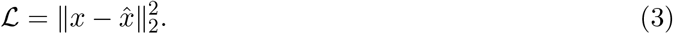

#### Linear probes

To assess the interpretability of latent dimensions, we use linear probing^19^ to identify components predictive of pathogenicity. We apply this to a dataset of single missense mutations with corresponding EVE scores. For each mutation, we extract residue-level embeddings from layer 24 of the pretrained ESM2 model and mean-pool them across the sequence to obtain a fixed-length vector, which is then passed through the pretrained SAE. We fit a ridge regression model on the resulting SAE embeddings to predict EVE scores. The dataset is split 80%/20% into training and test sets, and we perform 5-fold cross-validation on the training set to select the optimal regularization parameter *α* ∈ [0.001, 0.01, 0.1, 1, 10, 100]. The final model achieves a Spearman correlation of *ρ* = 0.75 on the held-out test set.

To interpret the latent space, we identify the five latent dimensions most positively and negatively associated with pathogenicity. For visualization, we highlight representative dimensions from each group on the wild-type OLF sequence. Among the pathogenic-associated latents, we observe that most active residues are polar or aromatic, consistent with known biochemical drivers of dysfunction. A few residues, including F467 and A488, did not follow this trend and were excluded from Figure 3b to better illustrate the dominant signal.

### B Full Visualization of the Phylogenetic Tree

Figure 4 details a full phylogenetic tree of OLF homologs. The sequences collected from UniRef100 comprise 73 taxonomic groups, encoding evolutionary signals from a diverse array of sources.

**Figure 4:**
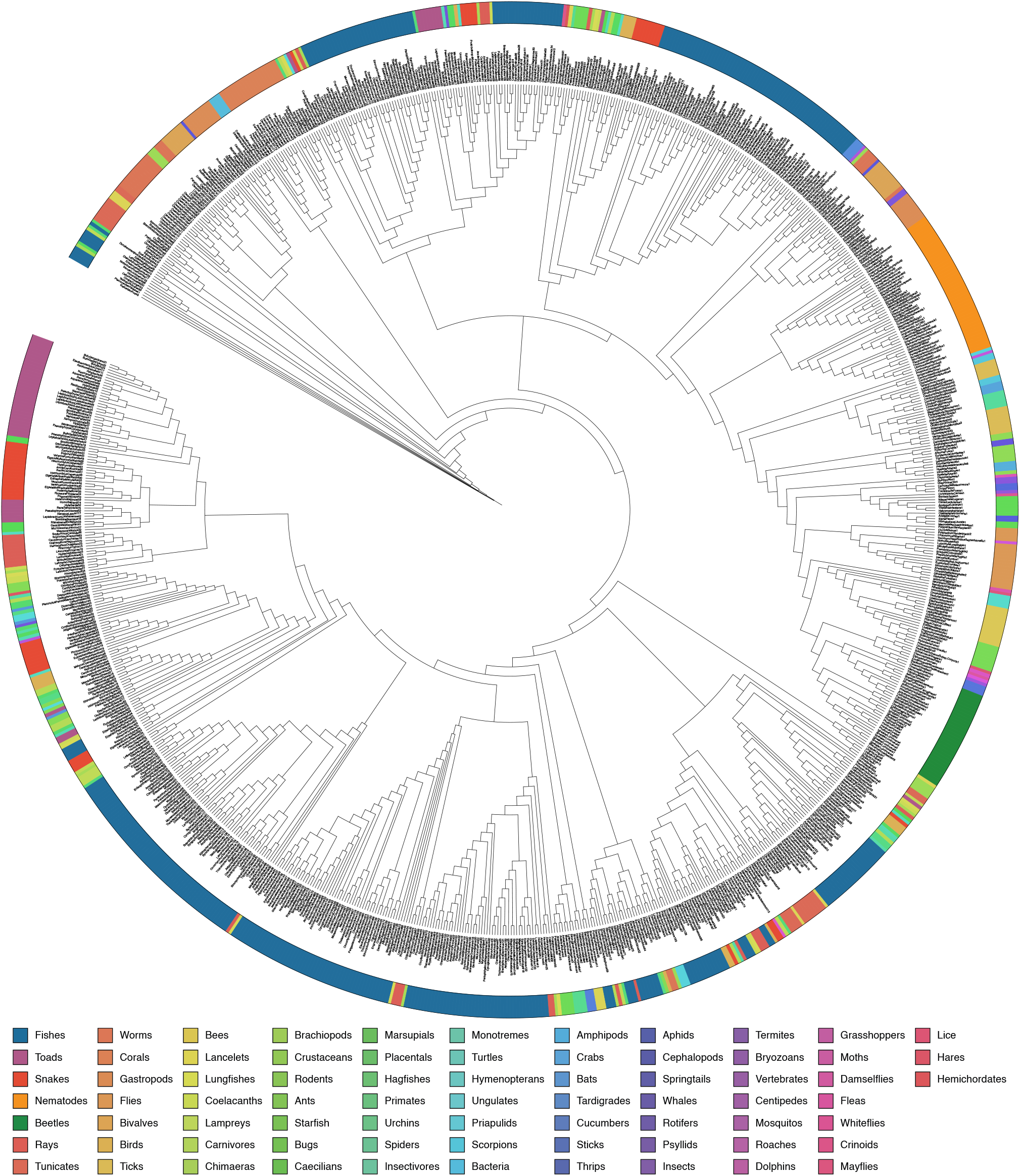
Phylogenetic tree of OLF homologs, collected from UniRef100.

### C Mutations

In this study, we assessed 32 mutations with known labels. The 7 benign mutations are as follows: Thr293Lys, Val329Met, Glu352Lys, Thr353Ile, Lys398Arg, Ala445Val, Lys500Arg. The pathogenic mutations comprise: Cys245Tyr, Gly246Arg, Gly252Arg, Arg272Gly, Glu300Lys, Glu323Lys, Gly364Val, Gly367Arg, Pro370Leu, Thr377Met, Thr377Lys, Asp380Ala, Lys423Glu, Val426Phe, Cys433Arg, Tyr437His, Ile477Asn, Ile477Ser, Tyr479His, Asn480Lys, Pro481Leu, Pro481Thr, Ile499Phe, Ile499Ser, Ser502Pro.

## Footnotes

1 GOLF is publicly available at https://github.com/amirgroup-codes/GOLF.git

